# Metagenomic insights into the development of microbial communities of straw and leaf composts

**DOI:** 10.1101/2024.07.08.602579

**Authors:** Anastasiia K. Kimeklis, Grigory V. Gladkov, Olga V. Orlova, Tatiana O. Lisina, Alexey M. Afonin, Tatiana S. Aksenova, Arina A. Kichko, Alla L. Lapidus, Evgeny V. Abakumov, Evgeny E. Andronov

**Author notes:** Address correspondence to Anastasiia K. Kimeklis,. Anastasiia K. Kimeklis and Grigory V. Gladkov contributed equally to this work. The authors order was determined based on seniority.

## Abstract

The objects of this study are microbial communities formed on two types of contrasting lignocellulosic biomass. To study them, a long-term experiment was set up and conducted on the decomposition of oat straw and leaf litter using soil as a microbial inoculum. Combined analyses of enzymatic activity and NGS data for 16S rRNA gene amplicon and full metagenome sequencing was applied to study taxonomic, CAZy (Carbohydrate-Active enzymes), and PULs (Polysaccharide Utilization Loci) composition of microbial communities at different stages of decomposition between substrates. using a combination of 16S rRNA gene amplicon and long-read nanopore sequencing. In straw degradation, the microbial community demonstrated higher amylase, protease, catalase, and cellulase activities, while peroxidase, invertase, and polyphenol oxidase were more active in leaf litter. Consistent with this, the metagenome analysis showed that the microbiome of straw compost was enriched in genes for metabolic pathways of simpler compounds. At the same time, there were more genes for aromatic compound degradation pathways in leaf litter compost. We identified 9 MAGs (Metagenome-assembled genomes) as the most promising prokaryotic decomposers due to their abnormally high quantity of PULs for their genome sizes, which were confirmed by 16S rRNA gene amplicon sequencing to constitute the bulk of the community at all stages of degradation. MAGs classified as Bacteroidota (*Chitinophaga*, *Ohtaekwangia*), and Actinomycetota (*Streptomyces*) were found in both substrates, while those from Bacillota (*Pristimantibacillus*) were specific for leaf litter. The most frequent PULs were specialized on xylans and pectins, but not cellulose, suggesting that PUL databases may be lacking PULs for complex substrates.

**Importance:** Our study explores the microbial communities from natural ecosystems, like soil and lignocellulosic waste, capable of decomposing lignocellulosic substrates. We used a comprehensive approach with chemical analyses of the substrates, amplicon and full-metagenome sequencing data. We have shown that such communities may be a source of identifying the highly effective decomposing species with novel PULs.

## Introduction

Most of the organic carbon storage in terrestrial ecosystems is located in the soil and plants, where it is involved in the global carbon cycle between living and dead biomass, atmosphere, and geosphere (1). Annually several hundred billion tons of lignocellulose biomass in the form of agricultural and forestry residues, food, and industrial wastes are produced (2, 3). Thus, its decomposition is a major part of organic carbon circulation in the terrestrial biomes (4). Recently, plant residue decomposition became the focus of metagenomic studies, but due to the high diversity of types of lignocellulosic substrates the process is also highly variable, and its specific features are still poorly understood (5). While the recirculation of plant litter in agricultural ecosystems receives increasing attention (6–8), the knowledge about biomass turnover in natural ecosystems remains limited (9). Yet, leaf litter, as a major component of the annual forest litterfall (10), and the primary component of soil formation, takes longer periods to decompose than straw, a part of agricultural waste, which is linked to the differences in their chemical composition (11, 12).

Lignocellulose biomass is a complex substrate, mainly consisting of recalcitrant cellulose, hemicellulose, and lignin in varying proportions (13, 14). Most of these compounds are degraded by enzymes, encoded by genes united in the CAZy database, including families of Glycoside Hydrolases (GHs), GlycosylTransferases (GTs), Polysaccharide Lyases (PLs), Carbohydrate Esterases (CEs), Auxiliary Activities (AAs), and Carbohydrate-Binding Modules (CBMs) (15). These genes are often combined in gene clusters, providing a full decomposition of a carbohydrate polymer (16). Historically, a cluster organization was shown for Bacteroidetes (17), where it was called Polysaccharide Utilization Loci (PUL), which represented the set of physically linked genes organized around a *sus*CD gene pair (18). Later cluster organization was shown for other phyla, not obligatory linked with these genes (19). So, for the automatic search of such structures, CAZyme gene clusters (CGC) were proposed, which encode at least one CAZyme, one transporter, one transcriptional regulator, and one signaling transduction protein (16). Still, while PULs are curated and experimentally accredited working gene clusters with known substrates, CGCs are not. So, PULs are highly sought after in natural ecosystems, but they are mostly described for gut microbiota (20). The most frequent carriers of PULs there belong to Bacteroidota and Bacillota (21). Whether PULs are limited to these environments and phyla is still not very clear.

In natural ecosystems, the initial sequestration of carbon is accomplished by soil microorganisms (22). Soils represent a global diversity storage of microbiota capable of decomposing lignocellulosic substrates. It is shown that communities from carbon-rich soils are more capable of assimilating organic matter (23). Thus, such soils can be used as a source of active microbiota. The soil of Chernevaya taiga (Tomsk, Russia) forms in the mid-mountainous area in the conditions of low eluviation of nutrients and low erosion (24). The unique combination of environmental factors leads to the gigantism of the perennial grass cover (25, 26), which annually leaves up to 400 g/m^2^ of lignocellulosic biomass to be processed by soil microorganisms. This results in the accumulation of the thick humic layer, sometimes reaching up to 70 cm in depth. The microbiota from this soil has great potential in decomposing cellulosic substrates, but this issue hasn’t been studied before. So, the goal of our study was to reveal this potential by setting up a long-term decomposing experiment, where the soil of Chernevaya taiga was enriched with two contrasting lignocellulosic substrates (straw and leaf litter) to reveal which parts of the microbial community would be more advanced in the new conditions and to reveal differences in the functional potential of these communities. We monitored this process by evaluating the shift of chemical composition of the substrates, taxonomic composition, and metagenome composition of the decomposing microbial communities with a focus on CAZy and PULs organization. The combination of Illumina sequencing of 16S rRNA gene and Oxford Nanopore metagenome sequencing was used to reveal the most probable active bacterial members of these communities.

## Materials and methods

### 1. Experiment setup

As a model of the decomposition of two contrasting in composition cellulosic substrates, we set up the experiment of composting oat straw and leaf litter with the soil-based inoculate.

The soil of Chernevaya taiga was collected near Tomsk, Russia in 2019. The forest is formed by a stable population of *Populus tremula* and *Abies sibirica*. The forbs are comprised of Siberian and Asia-specific tall-herbaceous perennials - *Euphorbia lutescens*, *Saussurea latifolia*, *Heracleum dissectum*, *and Alfredia cernua*. The soil type is Umbric Retisol, the coordinates are N 56.30693 E 85.47063. Its chemical characteristics and microbial composition were reported earlier (24, 25, 27, 28). Briefly, the topsoil was slightly acidic (pH = 5,8-6,2), with high organic carbon (2,4-9,9%) and nitrogen (0,2-0,6%) content. The phosphorus content reached 1000 mg/kg, comparable to the soils from tropical rainforests – one of the most productive ecosystems (29). To create a concentrated cellulolytic inoculum for the main experiment, a preliminary enrichment of the decomposing microbial community from soil was held in 2019-2020 for 14 months. For this 500 g of topsoil was mixed with 25 g (20:1 in mass, 1:1 in volume) of air-dry substrate (shredded oat straw or birch leaf litter) in 1-liter containers and incubated in a thermostat at 28±2 °C with the maintaining of 60% moisture content. After 8 months additional 5% of the substrates were mixed in. The resulting compost was used as an inoculate for the main experiment to accelerate the decomposition process.

For the experiment, two types of lignocellulosic biomass were used: (1) oat straw and (2) a mixture of fallen leaves of birch, oak, aspen, and willow (“leaf litter”). The chemical characteristics of the substrates are given in Table A1. Before setting the experiment, the substrates were shredded into particles 1-3 cm in size. For each type of lignocellulosic biomass 3 kg of air-dry substrate, 300 g of compost inoculum, and 9 L of water were mixed into a polypropylene 60-liter tub. Nine nylon bags with 5 g of air-dry substrate in each were placed inside the mixed mass for subsequent assessment of mass loss during composting. The top surface of the composted substrates was covered with plastic film to prevent drying out. Composting was carried out at a constant temperature of 28±2°C in a thermostat room. Moisture content was controlled once a week, and the substance was stirred once every two weeks.

Sampling was carried out three times during the experiment based on the external signs of the decomposition stage, which differed for the two substrates: straw on days 9, 61, and 111 and leaf litter on days 9, 111, and 174, in total six samples. Further, these dates will be referred to as the early phase (day 9 for straw and leaf litter), middle phase (day 61 for straw and day 111 for leaf litter), and late phase (day 111 for straw and day 174 for leaf litter). The chemical analyses were performed right after collecting, while samples for enzymatic and molecular analyses were stored at −20°C until the end of the experiment.

### 2. Sample processing

As a measure of a decomposition process, we evaluated mass loss of the substrate, respiration rates, chemical composition, and enzyme activity for six collected samples. The amount of the decomposed substrate was quantified by measuring the dry weight of the contents of nylon bags at the time of sample collection. The activity of the microbial biomass was assessed through the respiration of the substrate, expressed in the amount of the release of carbon dioxide using the alkali absorption method (30). Enzymatic analyses included amylase, catalase, exocellulase, endocellulase, peroxidase, polyphenoloxidase, invertase, and protease activity (31) which were carried out at the Agrophysical Research Institute. Other chemical analyses included the determination of pH, total carbon, nitrogen, cellulose, hemicellulose, and humic compound content (32, 33), which were conducted for the unaltered lignocellulosic substrates and samples of decomposed ones. The pH values were measured using pH meter F690 (Beckman Coulter, Inc., USA). The content of organic carbon (TOC) was expressed through the ash content in the substrates. The content of total nitrogen (TN) was measured on an automatic nitrogen analyzer (Boshu, Switzerland).

For the amplicon sequencing total DNA was extracted from the 6 experiment samples in triplicate using RIAM protocol (34). Pair-ended libraries were prepared using primers F515-GTGCCAGCMGCCGCGGTAA-3’/R806-GGACTACVSGGGTATCTAAT (35) for 16S rRNA gene and ITS3-GCATCGATGAAGAACGCAGC/ITS4-TCCTCCGCTTATTGATATGC (36) for the ITS2 fragment and sequenced on the Illumina Miseq platform (Illumina, Inc., USA).

DNA for the metagenome sequencing was extracted from the same 6 experiment samples, pre-stirred in liquid nitrogen, using the NucleoSpin® Soil Kit (Macherey-Nagel GmbH & Co. KG, Germany). The metagenome libraries were prepared using the Ligation Sequencing Kit 1D (Oxford Nanopore Technologies, UK) and sequenced on the MinION platform with the Flow Cell 9.4.1 rev D (Oxford Nanopore Technologies, Oxford, UK).

### 3. Data analysis

The amplicon sequencing data was processed in R Studio (37) using the DADA2 pipeline (38) as described earlier (39), including analysis of alpha (observed and Shannon (40) indices) and beta-diversity (NMDS (41) with Bray-Curtis distances (42)). Taxonomic identification was performed using Silva 138.1 (for 16S rRNA gene) (43) and Unite (for ITS amplicons, general release 02.02.2019) (44) databases. The Canonical Correspondence Analysis (CCA) (45) was performed to connect the microbiome composition of the composting substrates with the variability of their chemical characteristics. The data analyses of amplicon sequencing and analysis of chemical characteristics were performed using vegan (46), phyloseq (47), and ANCOM-BC (48) packages.

Metagenomes of the microbial decomposer communities of straw and leaf litter were basecalled by Guppy (49), assembled with Flye 2.9 using modifiers -meta and -nano-raw (50), and polished with medaka (51). The program kraken2 (52) in combination with the taxonkit utility (53) was used to determine the taxonomic composition of the metagenomes. The GTDB 214 database (54) was used as a reference base for prokaryotes (using the kraken2 struo2 database creation utility), and the PlusPF database based on the RefSeq NCBI database was used to identify eukaryotes. The obtained results were combined with the GTDB database being prioritized for prokaryote taxonomy divergence. Functional annotation of the metagenome was performed using EggNOG-mapperv2 (55) via Diamond with the --frameshift parameter (56). The differences in gene composition between metagenomes of different substrates were assessed using differential expression analysis in limma/voom from edgeR library (57). The annotation results were normalized by the sequencing depth, acquired by minimap2 (58) and samtools (59), and contigs with length less than 10000 were filtered out. Significantly different KOs between substrates were further processed by MinPath (60) program. DRAM annotator data was used to select genes responsible for lignocellulosic substrates decomposition based on functional annotation.

MAGs were isolated using the Semibin2 software (61). After polishing of the obtained draft bins by the medaka program, the bins have been selected for quality (medium (62) and high quality (63)) by the CheckM2 program. Taxonomic annotation was performed by gtdbtk (64). PUL search in MAGs was performed using dbCAN in dbCAN-PUL (65). The rRNA genes were isolated from the bins by the barrnap program (66) and aligned to the amplicon sequencing results. The results were visualized in the R software environment.

The packages versions and scripts used in the study are available at the repository https://github.com/crabron/clusters.

## Results

### 1. Chemical characteristics of the composting process

The decomposition process was evidenced by a decrease in respiration, and the loss of substrate mass, carbon, cellulose, and hemicellulose content, with differing dynamics for two substrates (Figure 1A). During the oat straw decomposition, the microbial activity (accessed by the carbon dioxide emission from substrates) was the highest in the early and middle phases and rapidly declined towards the end, indicating the completion of the decomposition of the most available organic compounds. The leaf litter rates in the early and late phases were not significantly different, which may be linked to the initial smaller content of available nutrients. Coinciding with the microbial activity, the process of decomposition for both substrates was marked by the mass loss of the decomposing substrate, which coincides with the loss of organic compounds. For the oat straw, 72.65% of the mass was decomposed by the late phase of the experiment, whereas the leaf mass loss by the late phase amounted to only 47.55%. In both substrates the hemicellulose undergoes decomposition by the early phase, while the cellulose content in the substrate remains practically unchanged. A significant decrease in cellulose content was detected in the middle phase for the oat straw and the late phase for the leaf litter. This also could be linked to a higher content of humic compounds in straw compared to leaf compost.

**Figure 1.**
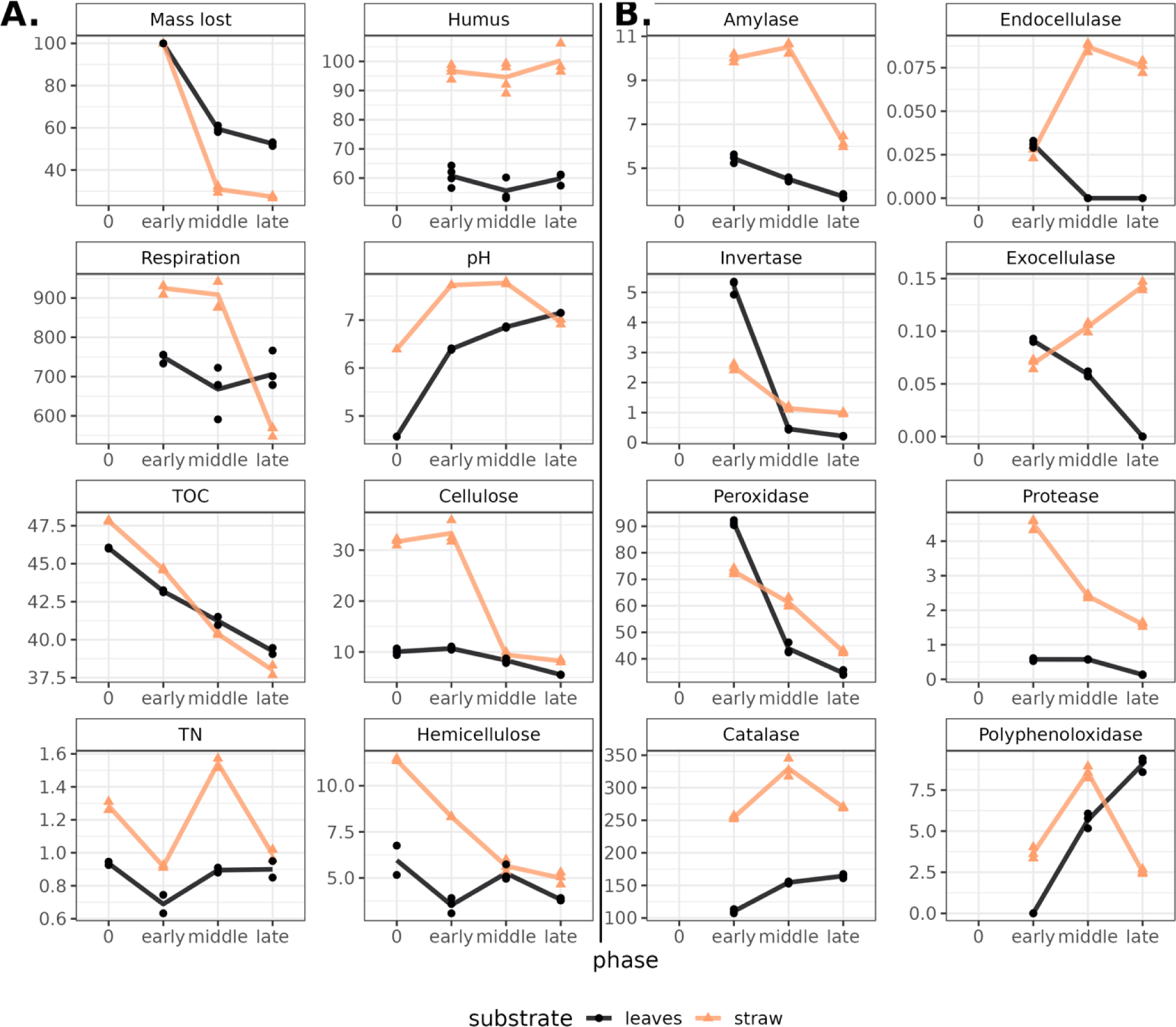
(A) Chemical characteristics and (B) enzyme activity in composts (leaves – black line, straw - yellow) during decomposition. Phase “0” stands for the characteristics of the cellulosic substrate without the soil-based inoculum. The Y axis corresponds with the absolute values of the parameters (See Table A1). Dots show replications (n=3) within samples. Lines connect the means. TOC – total organic carbon, TN – total nitrogen.

The total carbon content decreased gradually in both substrates as they decomposed, while the total nitrogen content shifted in waves: an increase followed by a decrease, indicating succession in the microbial communities. The leaf compost at the beginning of composting had a pH value slightly more acidic than the straw compost due to differences in substrate acidity. During the composting process, an increase in pH was observed for leaf litter and a slight decrease for straw.

The dynamics of the enzyme groups of interest were shown to shift according to the stage of degradation and substrate (Figure 1B). Both oat straw and leaf litter were characterized by a decrease in the activity of peroxidase, invertase, amylase, and protease by the late stages of decomposition. Agreeing with the data of chemical analysis, higher enzyme activity was shown in the decomposition of oat straw. Straw compost showed higher activity of catalase, endo-, exocellulases, peroxidases, and proteases. Leaf litter was more active in invertase. Polyphenoloxidase in straw compost was the most active in the middle phase, while in leaf litter it was in the last. Thus, significant differences in the rate and efficiency were observed in the decomposition of cellulose-containing substrates differing in nutrient availability.

### 2. Taxonomic composition of the composts

After filtering unidentified to phyla ASVs and organelle reads were obtained total 2244 phylotypes were identified for the 16S rRNA gene libraries and 250 for the ITS2 fragment. We observed a different pattern in taxonomic dynamics for prokaryotes and eukaryotes of microbial communities during plant residue decomposition. For prokaryotes, there was an increase in richness (number of observed ASV) of both composting substrates from early to late stages (Fig. 2A). Evenness (inverted Simson index) increased in leaf litter compost and decreased in straw compost. Both richness and evenness of the eukaryotic component were characterized by a decline in diversity from the early to the late decomposition stages. The richness values were different in the beginning (in leaf litter higher), but at the late stage, they were equally low. Initially, the evenness values were similar, but in straw, at the later stages, values were higher. Thus, the prokaryotic and eukaryotic components of the decomposing community were characterized by reverse dynamics - an increase in the richness of the former and a decrease in the latter.

**Figure 2.**
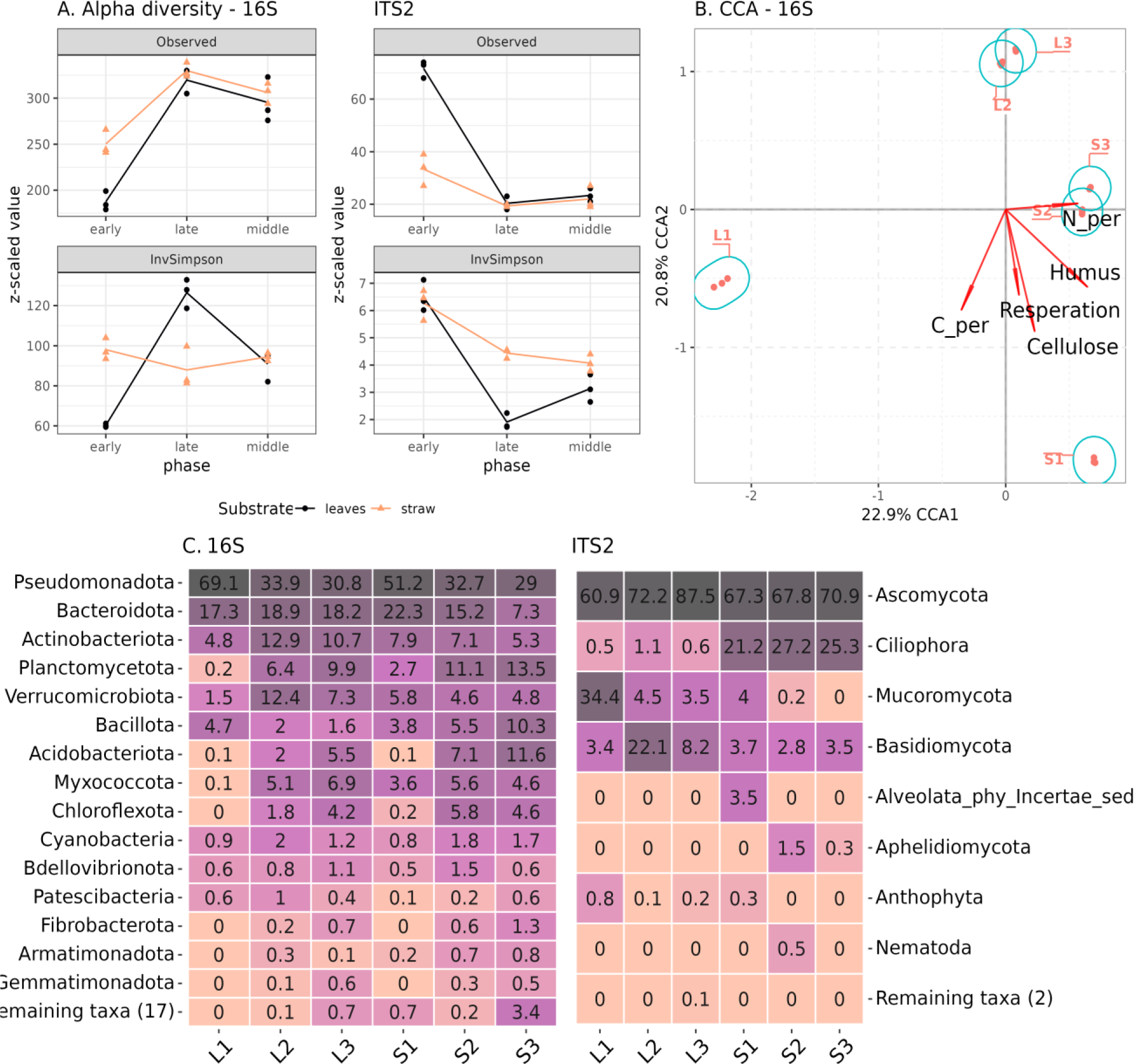
Prokaryotic and eukaryotic composition of composts accessed by 16S rRNA gene and ITS2 fragment Illumina sequencing. (A) Alpha diversity, (B) CCA for 16S rRNA, (C) Phylum composition (relative abundance). S1-S3 – samples from straw compost, L1-L3 – from leaf litter.

The beta-diversity of the microbiomes showed that the prokaryotic communities from the early phases of both composts were the most distinct from each other, while middle and late phases for each substrate were similar (Fig. 2B). The same was observed for eukaryotic communities, except that all phases of straw compost were quite close in composition (Fig. A1). According to the CCA, carbon content coincided with the dynamics of the decomposition, while respiration, pH, TN, cellulose, and humus content were associated with the substrate differences.

Since the experiment used a soil-based microbial inoculum for the initiation of the decomposition, the taxonomic composition of its prokaryotic community was already very diverse at the phylum level. The early phase consisted mostly of Pseudomonadota and Bacteroidota phyla, but during the composting process their relative representation decreased in favor of other phyla (Acidobacteriota, Planctomycetota, Myxococcota, Chloroflexota, Cyanobacteria) (Fig. 2C). In the straw compost Bacillota increased their abundance during decomposition, while in leaf litter – decreased. On the genus level in the early phase, both composts were dominated by *Flavobacterium*, *Pseudomonas*, *Pseudoxanthomonas*, and *Chitinophaga* (Figure A2). Additionally, in straw compost we detected *Devosia*, *Luteimonas*, and *Sphingobacterium*; in leaf – *Paenibacillus*, *Novosphingobium*, *Allorhizobium*, and *Galbitalea*. In the later phases, both composts were inhabited by *Ohtaekwangia*, *Acidibacter*, and *Steroidobacter*. Straw was characteristic of *Bacillus*, *Sphaerisporangium*, and *Clostridium*; leaf – *Terrimonas*, *Verrucomicrobium*, and *Bauldia*. The diverse eukaryotic community of the early phases of decomposition was gradually replaced by *Ovatospora* (Ascomycota) in both composts, which was especially characteristic of leaf litter (Figure 2C, A2). The late phase of the straw compost became inhabited by ciliates *Gastrostyla* and *Gonostomum*. Potentially pathogenic fungi *Alternaria*, *Pyrenophora*, *and Lichtheimia* were present at the early phases of decomposition but stopped detecting in the mature composts.

### 3. Functional composition of the composting substrates

After Nanopore sequencing, six metagenomes (three for straw and three for leaf litter) with a total read length of 190.4 Gbase were obtained. Following assembly and subsequent annotation, 14691 ko categories were identified in them. These were filtered and 5793 were used for the differential expression analysis. CAZy composition, attributed to polysaccharides utilization, didn’t show significant variability between substrate types or phases of decomposition, except for the early phase of leaf compost, which had the lowest values across all samples and DRAM categories (Figure 3A). The most abundant CAZy gene categories were amorphous cellulose, mixed-linkage glucans, xylans, xyloglucan, and chitin.

**Figure 3.**
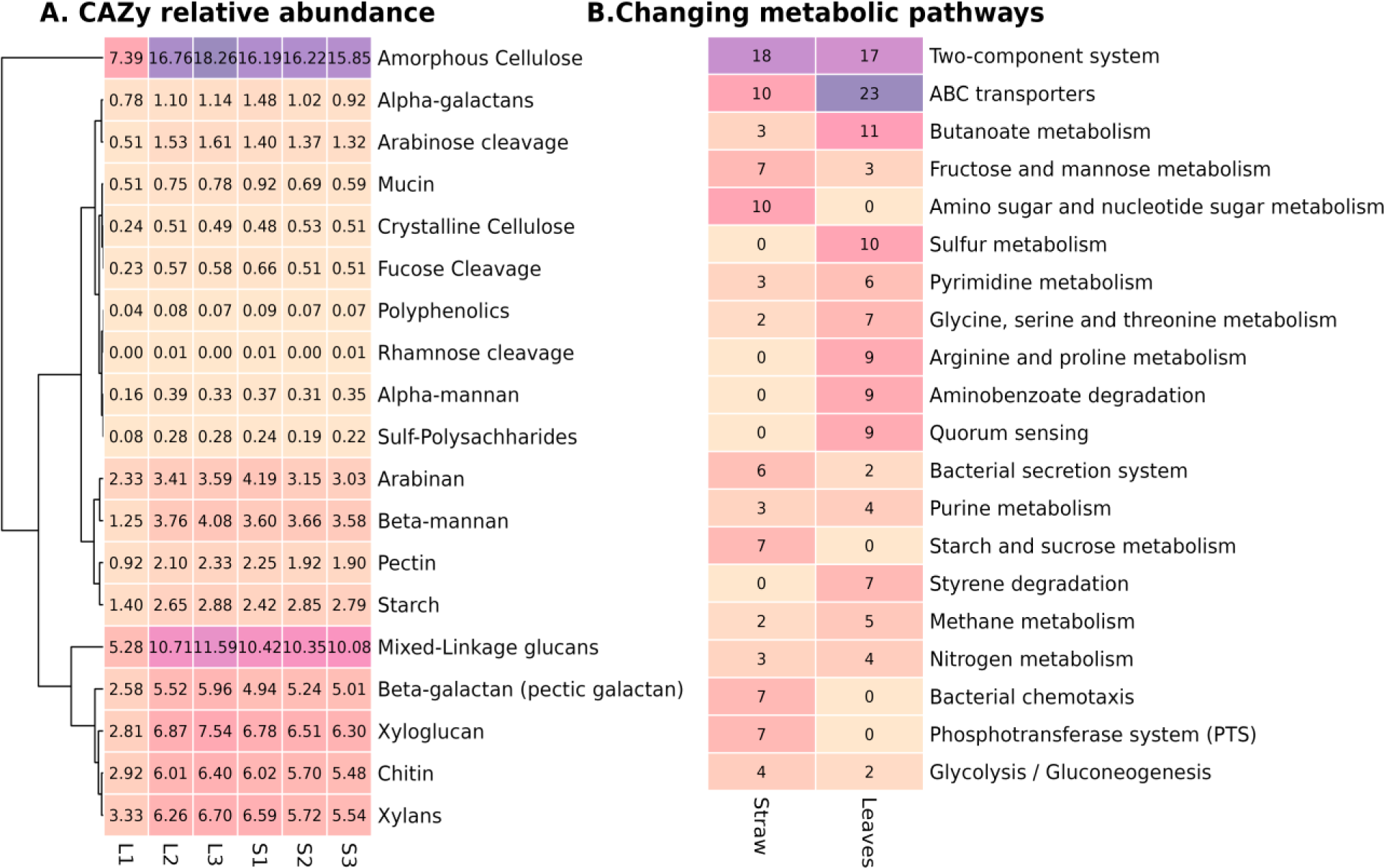
Metagenome characteristics of the contrasting composts: (A) Relative abundance (in percentiles from the ORFs from the metagenome) of CAZY genes united in DRAM categories in the six samples, (B) Number of KO (KEGG orthology) categories in the 20 most abundant metabolic pathways, significantly different between compost types.

To access significant differences between substrates we performed an enrichment analysis. As a result, 263 KEGGs were significantly increased in straw and 258 – in leaf litter (Figure A3). The most represented category was the two-component system, each substrate was more abundant in specific KO from this category (Figure 3B). Differences in the two-component system suggest that the leaf compost microbiome may experience catabolite repression, low nitrogen availability, and oxygen limitation (Figure A4). The second most abundant category was ABC transporters, and the leaf compost had more than twice the amount of KO compared to straw. Notably, leaf litter compost was more abundant in taurine and alkanesulfonate uptake, which coincides with the fact that it was also more abundant in KO from sulfur metabolism. Straw compost was enriched in the metabolism of simpler compounds (starch and sucrose, amino sugar, fructose, and mannose), while leaf litter was more abundant in KO from pathways of aromatic compounds degradation (xylene, benzoate, aminobenzoate, furfural, dioxin, hydroxy phthalate) (Figure 3B, Table A2). The metagenome of the straw compost had more KO, indicating the presence of microbial interaction – bacterial chemotaxis, antibiotic resistance, and flagellar assembly. Both composts were enriched in nitrogen metabolism, but straw compost had a pronounced denitrification pathway. Diversification on secretion type systems was also noted – while straw compost was more prevalent in types II, III, and VI, leaf litter was in types I and IV. Straw was enriched in biotin and zinc uptake.

Consistent with the Illumina data, a considerable amount of contigs in the metagenomic assembly of early phases of composting were annotated as belonging to the Pseudomonadota phylum, which decreased in later phases (Figure A5). Yet, even in the early phases proportion of genes associated with cellulose degradation in this phylum in the metagenomes was lower in both substrates compared to other phyla, e.g. Actinobacteriota, Planctomycetota, Bacteroidota, and Bacillota (Figure A5). The most characteristic was the difference between the earliest phase and the rest for both substrates. Oat straw was characterized by a high proportion of endocellulases in Bacillota and Verrucomicrobiota representatives. Further, the representation of these gene groups in the minor communities only continues to increase. For the leaf litter community, lower diversity of these genes (cellulase, catalase, amylase) in minor phyla was observed, with higher representation in representatives of Actinobacteriota and Bacteroidota. So, during the decomposition, we observed an increase in taxonomic diversity of the functional genes.

### 4. PULs

A total of 57 high and 240 medium quality MAGs were obtained. Of these, 254 contained clusters of carbohydrate-active genes (CGCs). In turn, polysaccharide utilization loci (PULs) were identified in 188 genomes, of which 135 had clusters capable of cleaving the β-1,4 bond between glucose residues characteristic of cellulose and hemicellulose. We compared the CGC/PUL ratio for MAGs from different phyla (Figure A6) to elucidate possible distortions in the data we obtained, connected with the underrepresentation of non-Bacteroidota phyla in the dbCAN-PUL. All ratios were striving towards 50%, except MAGs from Acidobacteriota and Verrucomicrobiota, although it may be caused by a small number of CGCs in them. The completeness of the genome assemblies didn’t affect these results. Thus, PUL quantities should adequately describe cellulolytic potential in the observed MAGs.

The highest number of PULs was observed in representatives of Bacteroidota, Pseudomonadota, Actinomycetota, Bacillota, and Chloroflexota (Table A3). The most numerous substrates were pectin, xylan, and arabinan. Pseudomonadota were also abundant in PULs for substrates, not directly connected with cellulose degradation, like capsule polysaccharide synthesis, glycosaminoglycan, and starch.

The obtained data allow us to identify the most potentially active cellulolytic organisms from the microbial community. For this purpose, we have chosen 9 MAGs containing an abnormally high proportion of PULs relative to their genome size (Fig. 4A). These were representatives of Bacteroidota (*Chitinophaga*, *Ohtaekwangia*), Actynobacteriota (*Streptomyces*) and Paenibacillales (*Pristimantibacillus*). There were other bacteria, mostly belonging to the Pseudomonadota, which contained high amounts of PUL, but their pools used pectin as a predicted substrate or were involved in cell capsid degradation. We found a match with 5 phylotypes from 16S amplicon sequencing data (Fig. 4B). Two bacilli genomes belonged to the same phylotype, and one representative of *Chitinophaga* aligned to a minor phylotype probably due to nanopore sequencing errors. The remaining phylotypes belonged to the major component of the community. *Bacillus* and *Streptomycetes* were characteristic of the early stages of leaf decomposition, *and Chitinophaga* was most represented in the early stages of straw decomposition. The representative of *Ohtaekwangia* did not show substrate specificity in relative representation and appeared only at the late stages of decomposition.

**Figure 4.**
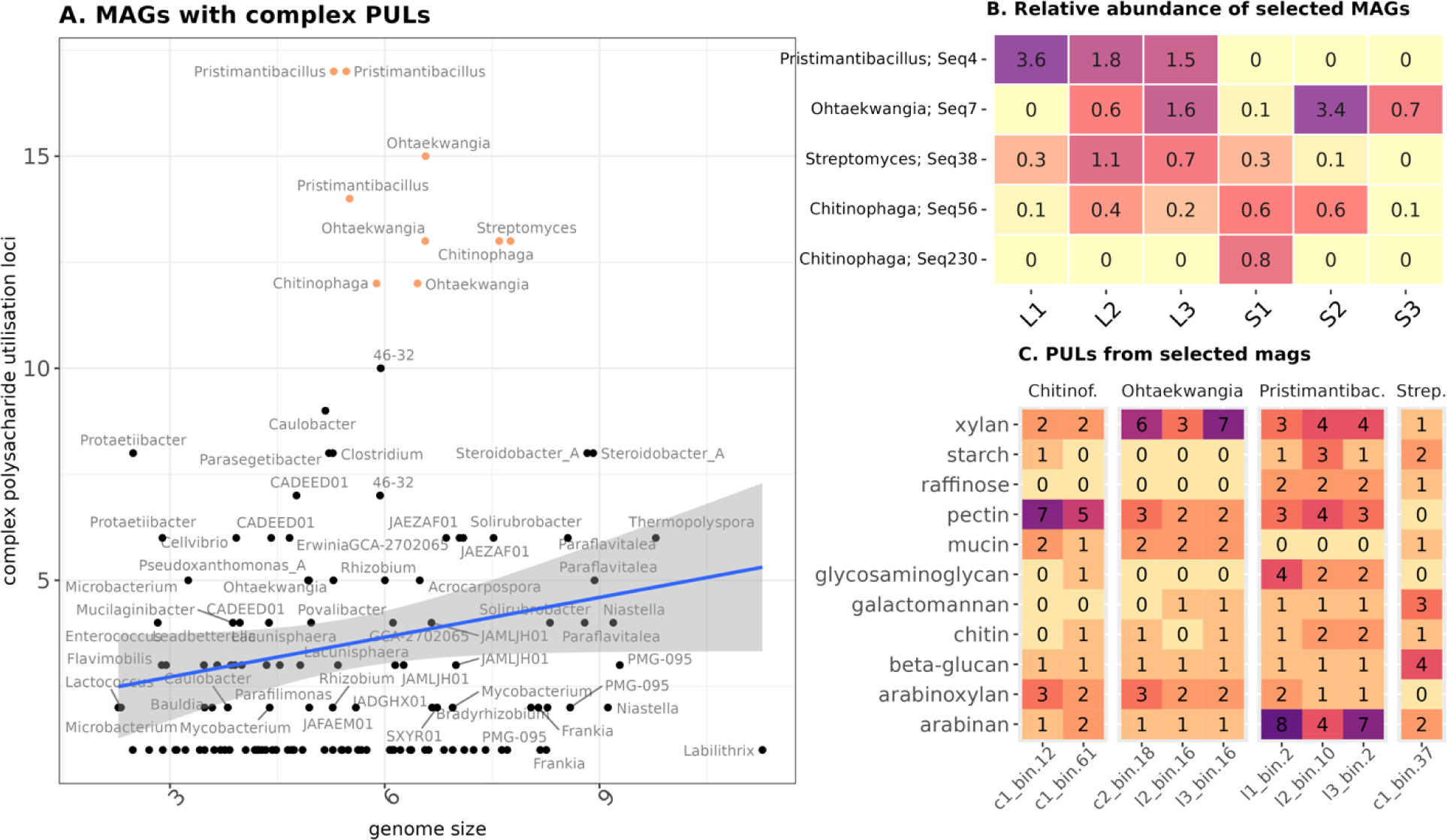
Characteristics of the most potential cellulolytic MAGs. (A) The ratio of PULs linked with lignocellulose degradation to the genome size. The group selected for further analysis is highlighted in color. (B) The presence of the chosen MAGs in the composts according to the Illumina data. (C) PUL substrates found in the selected MAGs. Genera names are given according to the GTDB annotation.

The identified PUL substrates varied between genera of MAGs. *Chitiniphaga* was the most abundant in PULs specific for pectin, *Ohtaekwangia* – xylan, *Pristimantibacillus* – arabinan, and *Streptomyces* – beta-glucan (Table A4). Other substrates, for which PULs were present, included arabinoxylan, mucin, glusaminoglycan, chitin, and starch. PULs of simpler polysaccharide compounds from MAGs assembled in this study shared high similarity with the reference ones – e.g. *Chitiniphaga* glucomannan, Streptomyces beta-glucan, *Ohtaekwangia* and *Pristimantibacullus* pectin (Figure A7). At the same time, PULs for xylans were mostly different in organization from the reference ones.

## Discussion

In the present study, cellulolytic communities of different decomposition stages of oat straw and leaf litter were selected as the objects of investigation. The use of a compost based on the soil of Chernevaya taiga soil as an inoculum was motivated by the fact that, although a significant number of active cellulolytic organisms have been described in the literature, most of them have been isolated from rumen and intestinal microbiomes (67), and soil microorganisms are extremely underrepresented in the databases. The advantage of soil microbiota over rumen is in the potentially higher adaptation to the decomposition of a wider range of plant residue types (68). Understanding which part of the microbial community has the greatest potential for lignocellulosic complex decomposition may allow a shift from microbial preparations based on individual strains to more effective ones based on microbial consortia (69, 70). This is specifically important because soil communities are highly diverse and relatively stable, which complicates the integration of microbial preparations based on single strains (71, 72). The soil of Chernevaya taiga, formed in unique environmental and geochemical conditions, is specifically promising in finding such microbiota.

We used two contrasting lignocellulosic substrates - oat straw and leaf litter. The dynamics of chemical characteristics and differences in the activity of the enzymes in both composts may indicate that oat straw in the experiment was more saturated with water-soluble hydrocarbons and proteins relative to the leaf litter, while the latter was more recalcitrant due to higher content of resilient materials, not covered by our analyses. Besides, while organic carbon content was similar, nitrogen content was higher in straw, so the C/N ratio, which affects the activity of microbiota (73, 74), was more favorable in straw than in leaf litter. The lower C/N ratio could also explain the release of mineral nitrogen in the middle phase of oat straw decomposition (75), which additionally facilitated the endo- and exo-cellulolytic activity of the microbiota. The presence of bacterivorous ciliates in straw compost may serve as further evidence of the faster turnover of the organic material (76). Consistent with this, in our experiment straw turned out to be a more accessible substrate and decomposed much faster than leaf litter.

According to the above, while both straw and leaf litter are cellulosic substrates, their resistance to decomposition is different due to chemical characteristics. This establishes conditions for the diversification of the initial microbial soil community. We saw this effect in the taxonomic structure of the compost microbial communities, which shifted significantly between the types of substrates. However, both composts shared similar patterns of microbiome development. Our choice of longer periods of sample incubation had a significant effect on the outcome: we detected long-term shifts in microbial and functional composition, with an increase in bacterial diversity and a decrease in fungi, coinciding with one of our previous experiments (39). The observations of dynamics at the taxonomic level are consistent with the earlier findings, specifically the prevalence of Pseudomonadota during the initial stages of straw colonization from soil and the diversification of the community in the later phases (8, 39). Genera detected by Illumina sequencing in both types of composts – *Pseudomonas*, *Bacillus*, *Flavobacterium, Sphingobacterium, and Novosphingobium* – were reported multiple times to contain potentially active cellulolytic species (77–80). Cellulose-degrading strains from *Pseudoxanthomonas suwonensis* (81) and *Luteimonas* (82) were isolated from the soil. So, while both composts were diversified in major microbiota, most of it was comprised of potential decomposers. Despite this, on the taxonomy level of analysis, we cannot determine which members of the community have this predisposition, especially considering that the microbiomes of the middle and late phases in both composts were very close, but the enzymatic activity between these phases shifted very prominently. Besides, earlier findings showed that only a minority of a bacterial community in the compost may be associated with lignocellulosic substrate degradation (6). So, we used metagenomic data to look for the potential active decomposers in both communities.

Despite the strong variation in the activity of cellulolytic-related enzymes between phases of decomposition and substrate types, we didn’t detect significant shifts in CAZy categories between the six analyzed metagenomes. Consistent with this, no enrichment of any CAZy families was shown between metagenomes of microbial communities isolated from different substrates (83). The activity of catalytic enzymes decreased by the late phases of decomposition, while the relative content of genes associated with lignocellulose decomposition remains stable. This can be attributed to the activity of their gene expression, which in turn is regulated at the cellular level by regulatory systems (e.g., two-component systems) responding to the substrate representation in the environment. This assumption is evidenced by the overrepresentation of diverse KO groups from two-component metabolic pathways between compost types. Another important observation is that the taxonomic diversity of genes associated with lignocellulosic complex decomposition between phases grew more actively than for other genes. This shows a greater involvement of minor phyla in cellulose degradation. This effect is particularly evident in the late stages of decomposition and is substrate dependent. A similar effect was shown previously, as we detected that a significant part of microbial taxonomic diversity at the later phases of straw colonization does not participate in cellulose degradation because it does not have genes for the corresponding enzymes in its genomes, while these groups of microorganisms can make a serious contribution to the community by shifting the representation of effective cellulolytic microorganisms (39).

To identify the most probable active decomposers of the microbial communities of the composts, we searched for MAGs with an abnormally high content of PULs relative to their genome size. The previous study on lignocellulose biomass decomposition indicated that the most promising degraders, according to the GH gene content, belong to Bacteroidota and Bacillota (5). Consistent with their findings, according to Illumina sequencing Bacteroidota and Bacillota phyla were abundant in both composts, and out of 9 MAGs, which we selected as promising active decomposers according to our criteria, 5 belonged to the first and 3 to the latter. Notably, Actinobacteriota were also reported to play an important role in the lignocellulose degradation (84), only one potentially cellulolytic MAG from this phylum was assembled, which may be linked to its genome size (85). All these MAGs were the representatives of both minor and major microbiota, indicating the involvement of the diverse ecological groups in the process of decomposition.

This data couldn’t be checked for significance, but the PULs specificity seems to be connected to the MAG taxonomic attribution, not the phase of decomposition, from which it was detected. Since we observed the taxonomic differentiation of potentially active MAGs between composts, we can assume that PUL specificity is also highly linked with the substrate type.

Interestingly, the study of Huang et. al (5) operated with CGCs and not verified PULs, because most of their data was not indexed in dbCAN-PUL. We managed to work on the PUL level, but most of their substrates were compounds, accompanying cellulose fibrils. Along with that, we the PULs, specific for cellulose degradation, were absent in almost all MAGs, selected as potentially active degraders. The PUL database mostly covers compounds, usually found in the gastrointestinal tract, which is the most frequent object for studying polysaccharide degradation.

All of the above shows that the design of effective lignocellulosic communities requires an integrated approach, which is not limited only to the analysis of individual enzyme groups or individual taxa. It is important to understand the principles of formation and functioning of cellulolytic microbial consortium to apply this knowledge to the formulation of highly effective microbial preparations.

## Conclusion

The novelty of our research lies in the development of new approaches to the search for promising cellulolytic microorganisms based on next-generation sequencing data. The composts based on two types of lignocellulosic biomass were studied using the combination of analyses of enzyme activities, Illumina Miseq read-based sequencing of 16S rRNA gene amplicon libraries, and metagenome analysis using Oxford Nanopore MinION long read technology to search for promising cellulolytic prokaryotes. The complementarity and convergence of the two sequencing methods in the context of soil metagenomics were demonstrated. Substrate specificity was found to be rather associated not with the presence or absence of certain CAZy gene families, but with their organization into gene clusters - PULs. The analysis of the genomes of potential cellulolytic microorganisms showed that some of them are substrate-specific and are major representatives of the microbial community. Historically PULs were found in the gut communities, but our work demonstrates that communities decomposing lignocellulose can also be a valuable source for their identification.

## Data Availability

Data are available at the NCBI BioProject ID PRJNA1131150 (http://www.ncbi.nlm.nih.gov/bioproject/1131150).

## Acknowledgments

We thank the Centre for Genomic Technologies, Proteomics and Cell Biology (ARRIAM, Russia) for performing the preparation and sequencing of gene and genomic libraries and the Agrophysical Research Institute for the enzymatic analysis. We also want to thank Sergei V. Loiko and Georgy I. Istigechev for organizing the field research of soil from the Chernevaya taiga under the leadership of Lapidus A.L.

## Author Contributions

Conceptualization, A.K.K. and G.V.G.; Data curation, G.V.G.; Formal analysis, G.V.G., O.V.O., and A.M.A.; Funding acquisition, E.E.A.; Investigation, A.K.K. and G.V.G.; Methodology, G.V.G., O.V.O., T.O.L., A.A.K., and T.S.A.; Project administration, E.E.A.; Resources, E.V.A, A.L.L., T.S.A., A.A.K., and A.G.P.; Software, G.V.G. and A.M.A.; Supervision, E.E.A.; Validation, A.K.K., G.V.G., and A.M.A.; Visualization, A.K.K. and G.V.G.; Writing—original draft, A.K.K. and G.V.G.; Writing—review & editing, A.K.K., G.V.G., A.M.A., and A.L.L. All authors have read and agreed to the published version of the manuscript.

## Institutional Review Board Statement

Not applicable.

## Informed Consent Statement

Not applicable.

